# Ethanol exposure alters Alzheimer’s-related pathology, behavior, and metabolism in APP/PS1 mice

**DOI:** 10.1101/2022.02.18.481066

**Authors:** Stephen C. Gironda, Stephen M. Day, Caitlin W. Clarke, J. Andy Snipes, Noelle I. Nicol, Hana Kamran, Warner Vaughn, Shannon L. Macauley, Jeff L. Weiner

## Abstract

Chronic ethanol exposure can increase amyloid-β (Aβ) and tau in rodent models of Alzheimer’s-disease (AD)-like pathology, yet the underlying mechanisms are poorly understood. In this study, a moderate two-bottle choice drinking paradigm was used to identify how chronic ethanol exposure alters Aβ-related pathology, metabolism, and behavior. Complementary in vivo microdialysis experiments were used to measure how acute ethanol directly modulates Aβ in the hippocampal interstitial fluid (ISF). Ethanol-exposed APPswe/PSEN1dE9 (APP/PS1) mice showed increased brain atrophy and an increased number of amyloid plaques. Further analysis revealed that ethanol exposure led to a shift in the distribution of plaque size in the cortex and hippocampus. Ethanol-exposed mice developed a greater number of smaller plaques, potentially setting the stage for increased plaque proliferation in later life. Ethanol also induced changes in N-methyl-D-aspartate and γ-aminobutyric acid type-A receptor (NMDAR and GABA_A_R, respectively) expression, possibly reflecting changes in the excitatory and inhibitory (E/I) balance in the brain. Ethanol exposure also led to a diurnal shift in feeding behavior which was associated with changes in glucose homeostasis and glucose intolerance. Ethanol exposure also exacerbated alterations in the open-field test and deficits in nest-building behaviors in APP/PS1mice. Lastly, an acute dose of ethanol bidirectionally altered hippocampal ISF Aβ levels – decreasing during the initial exposure and increasing during withdrawal. Acute ethanol exposure increased hippocampal ISF glucose levels, suggesting changes in cerebral glucose metabolism occur in response to ethanol. These experiments indicate that ethanol exacerbates an AD-like phenotype by altering Aβ deposition, behavior, and metabolism. Here, even a moderate drinking paradigm culminates in an interaction between alcohol use and AD-related phenotypes with a potentiation of AD-related pathology, behavioral dysfunction, and metabolic impairment.

**Highlights:**

- Chronic ethanol exposure increases brain atrophy in APP/PS1 mice.
- Chronic ethanol exposure increased the number of plaques in the brains of APP/PS1 mice.
- Chronic ethanol exposure led to dysregulated metabolism in APP/PS1 mice.
- Chronic ethanol exposure altered anxiety- and dementia-related behaviors in APP/PS1 mice.
- Acute ethanol exposure bidirectionally alters interstitial fluid (ISF) levels of amyloid-β in APP/PS1 mice during exposure and withdrawal.

## Introduction

Alzheimer’s disease (AD) is responsible for 60-80% of dementia cases and is the most common form of dementia. In the US, there are approximately 6 million people diagnosed with AD, which is expected to increase to 14 million by 2050 (Long and Holtzman 2019). According to the A/T/N framework, AD pathology is characterized by the aggregation of extracellular amyloid-β (Aβ) into amyloid plaques, the intracellular accumulation of tau into neurofibrillary tangles, and neurodegeneration (Jack, Bennett et al. 2016). Aβ aggregation and other pathological events precede the onset of cognitive decline and clinical diagnosis by ∼10-20 years (Jack, Knopman et al. 2010), making it important to identify risk factors that accelerate the onset, hasten the development, or exacerbate the pathological changes in AD. Epidemiological studies identified alcohol use disorder (AUD) as a significant risk factor for AD (Harwood, Kalechstein et al. 2010, Xu, Wang et al. 2017, Schwarzinger, Pollock et al. 2018, Zhornitsky, Chaudhary et al. 2021), yet there is conflicting evidence on how alcohol use promotes AD pathogenesis. On the other hand, some studies suggest that lower levels of alcohol (ethanol) consumption may reduce AD risk (Rehm, Hasan et al. 2019). Preclinical studies show that chronic administration of low, moderate, and high levels of ethanol can affect amyloid plaque pathology and amyloidogenic processing of amyloid precursor protein (APP) in multiple models of Aβ overexpression (Huang, Yu et al. 2018, Hoffman, Faccidomo et al. 2019). Unfortunately, differences in methodology make the results difficult to interpret. Therefore, questions remain as to whether ethanol exposure directly modulates Aβ levels, resulting in amyloid plaque pathology and its associated behavioral and metabolic deficits.

Recognizing this significant gap in knowledge and the critical need to better understand AUD as a risk factor for developing AD, we investigated how chronic ethanol exposure alters behavioral and metabolic disturbances associated with AD pathogenesis. Here, a well-validated mouse model of Alzheimer’s-related pathology and Aβ overexpression (APPswe/PSEN1dE9; APP/PS1) (Jankowsky, Fadale et al. 2004) was chronically exposed to moderate amounts of ethanol. The effects of ethanol on AD-related pathology, excitatory and inhibitory (E/I) receptors, metabolism, anxiety- and depression-related behaviors, and cognitive measures were then analyzed. In vivo microdialysis in APP/PS1 mice was used to measure how acute, systemic ethanol administration directly impacts Aβ levels and glucose metabolism in the interstitial fluid (ISF) of the hippocampus.

In this study, the long-term consumption of moderate amounts of ethanol via a two-bottle choice drinking paradigm induced changes in Aβ plaque number, plaque size, and brain atrophy. While moderate ethanol consumption did not directly alter APP levels or APP metabolism, it did lead to changes in glutamatergic and GABAergic receptor subunit expression. This suggests that ethanol altered neuronal activity, which is known to affect Aβ release and aggregation (Cirrito, Yamada et al. 2005, Cirrito, Kang et al. 2008, Bero 2011). Moderate ethanol consumption also led to disruptions in glucose homeostasis in both the CNS and periphery, which are known drivers of Aβ-related pathology (Macauley, Stanley et al. 2015, Stanley, Macauley et al. 2016). Moderate ethanol exposure also exacerbated behavioral deficits typically observed in APP/PS1 mice. The acute, systemic delivery of ethanol bidirectionally modulated Aβ levels in hippocampal ISF. Aβ levels decreased by nearly 20% during the initial exposure then increased by nearly 20% during withdrawal. This increase during withdrawal corresponded to the clearance of ethanol from the ISF. Collectively, this study provides evidence that ethanol exposure directly modulates Aβ levels. It lends additional support to the notion that chronic consumption of even moderate amounts of ethanol may exacerbate the development of AD-related pathology and behavioral deficits.

## Materials & Methods

### Animals

5.5 month-old male APPswe/PSEN1dE9 mice (Jankowsky, Fadale et al. 2004) (APP/PS1; The Jackson Laboratory; n=17) and age-matched wildtype B6C3 control mice (n=17) were used for the chronic drinking studies. All animals were housed individually in standard mouse cages under a 12-h artificial light–dark cycle. Room temperature and humidity were kept constant (temperature: 22 ± 1 °C; relative humidity: 55 ± 5%). Standard laboratory rodent chow and tap water were provided ad libitum throughout the experimental period. Mice underwent a battery of behavioral tests at baseline, and at various stages during ethanol exposure. A separate cohort of 3-month-old male APP/PS1 mice (n = 4-6) was used for acute ethanol exposure experiments. All experimental procedures were approved by the Committee on Animal Care and Use at Wake Forest School of Medicine.

### Experimental Design

At 4 months of age, APP/PS1 and control mice were run through a battery of behavioral tests to identify baseline differences in anxiety- and depression-related behavior. Following completion of this behavioral battery at 5.5 months of age, APP/PS1 and control mice were randomly assigned to either an ethanol exposure or water alone condition. Ethanol exposure was provided via a 10 week two-bottle choice paradigm (Huynh, Arabian et al. 2019). Mice were weighed before and after each drinking session. Throughout the 10 week drinking period, mice were assessed for changes in anxiety and AD-related behaviors. All behavioral assays were conducted during the three-day abstinence period. At the end of the study, mice were euthanized within 24-48 hours after the final ethanol exposure period (see figure 1a for experimental timeline). Briefly, mice were anesthetized with an overdose of sodium pentobarbital and transcardially perfused with ice-cold 0.3% heparin in DPBS. Prior to perfusion ∼200 μL of blood was collected from the left ventricle and transferred into EDTA-coated tubes and kept on ice. Tubes were centrifuged at 3000g for 5 minutes at 4°C and plasma was removed then flash-frozen in dry ice and stored at −80°C. After perfusion brains were removed, weighed, then bisected. The left hemisphere was fixed in 4% paraformaldehyde at 4°C, while the right hemisphere was dissected and flash-frozen in dry ice.

**Figure 1:**
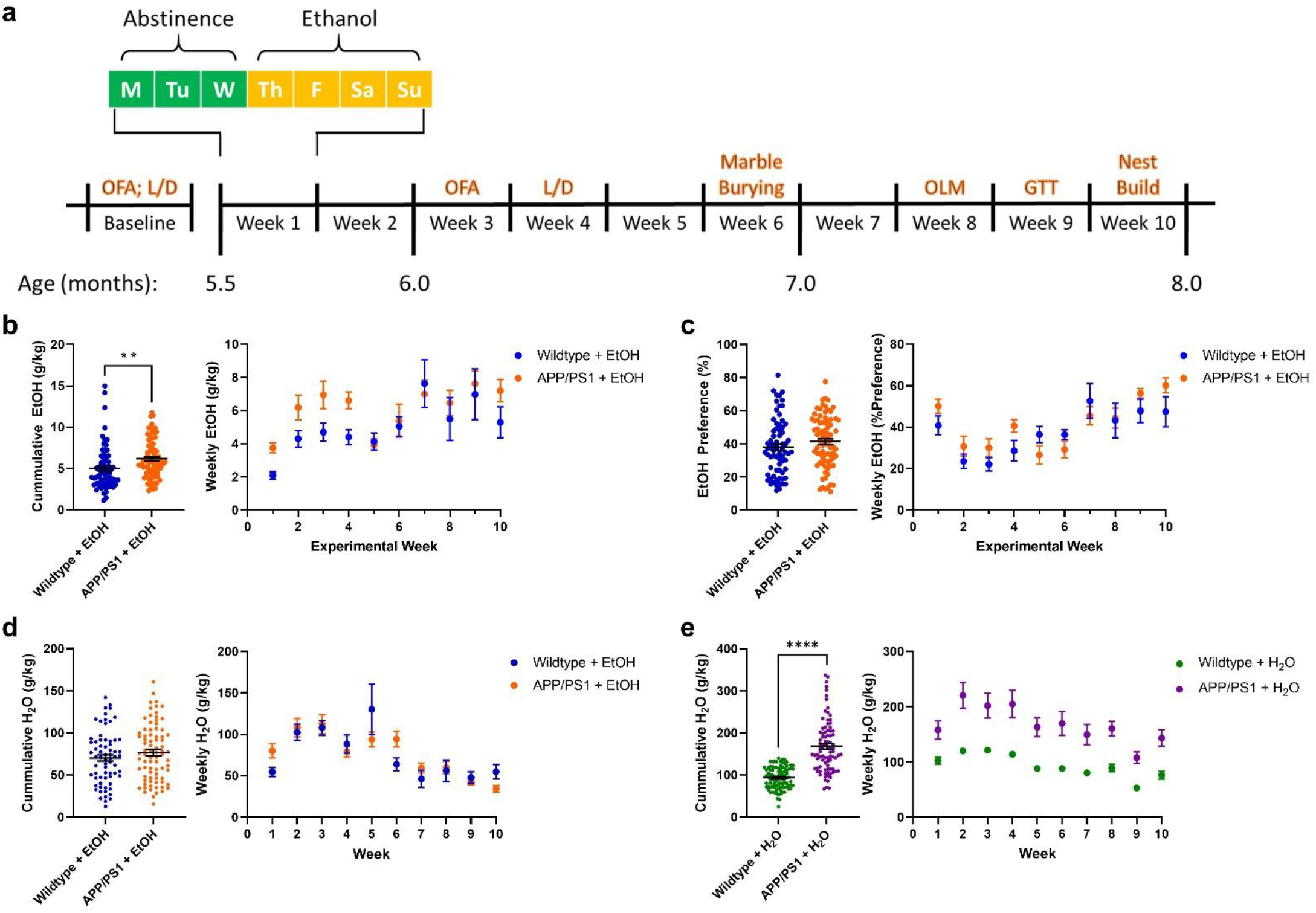
APP/PS1 mice consume more ethanol than control mice. a) Timeline for the experimental protocol. Full details are provided in the methods. b) Cumulative and average weekly EtOH intake (g/kg) from the 10-week plotted as a function of genotype. APP/PS1 mice consumed more EtOH than wildtype mice (p<0.01, unpaired t-test). c) Cumulative and average weekly EtOH preference (% total fluid) from the 10-week exposure period plotted as a function of genotype. No difference between EtOH preference between wildtype and APP/PS1 mice was observed (unpaired t-test). d) Cumulative and average weekly water consumption across the 10-week EtOH exposure in EtOH-treated mice. No difference in weekly water consumption was seen in EtOH-treated wildtype or APP/PS1 mice (unpaired t-test). e) Cumulative and average weekly water consumption across the 10-week EtOH exposure period in water-treated mice. APP/PS1 mice consumed more water than wildtype mice (p<0.0001, unpaired t-test). *p<0.05, **p<0.01, ****p<0.0001

### Two Bottle Choice Procedure

Mice were individually housed and exposed to a modified two bottle-choice drinking paradigm for 10 weeks (Huynh, Arabian et al. 2019). Briefly, home-cage water bottles were replaced with two 50 mL bottles containing H_2_O or a 20% ethanol solution. The H_2_O control group received only H_2_O in both bottles. Mice were given access to ethanol during their dark cycle for 12 hrs/day for 4 days/week, and the position of the bottles were alternated daily to avoid side preference. Mice were habituated to 20% EtOH over a two-week period by incrementally increasing EtOH concentration in the water from 5% to 20% (v/v). Fluid consumption was measured by weighing bottles before and after each drinking session. Ethanol is presented as g of ethanol per kg of mouse (g/kg). EtOH preference was calculated as a percent of ethanol intake to total liquid intake.

### Brain mass, Aβ immunohistochemistry, and X34 staining

Prior to sectioning, brains were cryoprotected in 30% sucrose then sectioned on a freezing microtome at 50 μm. Three serial sections (300 μm apart) through the anterior-posterior aspect of the hippocampus were immunostained for Aβ deposition using a biotinylated, HJ3.4 antibody (anti-Aβ_1-13_, a generous gift from Dr. David Holtzman, Washington University). Sections were developed using a Vectastain ABC kit and DAB reaction. For fibrillary plaques, free floating sections were permeabilized with 0.25% Triton X-100 and stained with 10 µM X-34 in 40% ethanol + 0.02M NaOH in PBS (Ulrich, Ulland et al. 2018). Brain sections were imaged using a NanoZoomer slide scanner and the percent area occupied by HJ3.4 or X34 was quantified using ImageJ software (National Institutes of Health) as previously described (Bero, Yan et al. 2011, Roh 2012). A histogram analysis was performed to quantify the frequency of each plaque by pixel size, excluding any plaques smaller than 10 pixels. Statistical significance was determined using a two-tailed, unpaired t-test (percent area) and a 2-way ANOVA with Šídák’s multiple comparisons post-hoc tests (size x frequency). Aβ deposition, amyloid plaque size, amyloid plaque, and neurofibrillary plaque area fraction were quantified by a blinded researcher. Data is represented by means ±SEM.

### Western blot

Western blot analysis was used to measure protein levels of APP processing enzymes and excitatory and inhibitory receptors. For APP processing enzymes, posterior cortical tissue was homogenized in 1X cell lysis buffer (Cell Signaling) supplemented with protease inhibitor cocktail (Roche), 1mM PMSF (Cell Signaling), 1mM DTT (Sigma-Aldritch), and a phosphatase inhibitor cocktail (Millipore) using a probe sonicator at 30% amplitude, 1 sec pulse with a 5 sec delay, 5 times while on ice. Tissue homogenates were then spun down at 10,000g for 10 minutes at 4°C and the supernatant was used for immunoblotting. Protein concentrations were analyzed using BCA protein assay kit (Pierce). For APP and APP c-terminal fragments (CTFs), 15 μg of protein were run in 15% tris-tricene gels to increase separation between CTF-β and CTF-α. All other proteins were run in 10% tris-trice gels. All gels were run using BioRad Protean mini then rapid-transferred to PVDF membranes using BioRad Semi-dry membranes (BioRad). Membranes were subsequently blocked using 5% BSA in 1X TBST for 1 hour and then incubated with primary antibodies overnight at 4°C. Secondary antibody conjugated with HRP-specific to primary antibody were incubated at room temperature for 1 hour in 1X TBST. The following primary and secondary antibodies were used for this study: APP (including CTFβ and CTFα; Invitrogen; CT695; 1:1000), BACE1 (Cell Signaling; 5606S; 1:1000), ADAM10 (Millipore; AB19026; 1:1000), IDE (Abcam; ab232216; 1:1000), GluN2A (Cell Signaling; 4025; 1:1000), GluN2B (Cell Signaling; 4212; 1:1000), GABAAR α5 (Santa Cruz; Sc393921; 1:1000), and β-actin (Millipore; MAB1501; 1:50,000), anti-mouse (Cell Signaling; 7076S; 1:5000), anti-rabbit (Cell Signaling; 7074S; 1:5000). Protein bands were visualized using chemiluminescence using ECL (EMD Millipore). Densitometric analysis was performed using ImageJ, and data were normalized to β-actin.

### qPCR

RNA isolation from mouse brain was performed as previously described (Musiek, Lim et al. 2013). Briefly, anterior cortex was homogenized by trituration through a 23-gauge needle in TRIzol (Invitrogen). Chloroform (1:5) was added then samples were mixed, and centrifuged (13,000g; 15 minutes; 4°C). Chloroform was removed, and samples were diluted 1:1 in 70% ethanol and purified using RNeasy columns and reagents (QIAGEN). RNA concentration was measured using a NanoDrop spectrophotometer. Reverse transcription was performed using a highcapacity RNA-cDNA kit (Applied Biosystems [ABI]) with 1 μg RNA per 20 μL reaction. Real-time qPCR was performed with ABI TaqMan primers and reagents on an ABI Prizm 7500 thermocycler according to the manufacturer’s instructions. Primers used: Grin2a (TaqMan; Mm00433802_m1), Grin2b (TaqMan; Mm00433820_m1), Gabra5 (IDT; Mm.PT.58.5845925), Actb (TaqMan; Mm01205647_g1). All mRNA measurements were normalized to Actb (β-actin) then to wildtype + H_2_O group mRNA levels.

### Synaptoneurosome preparation

Synaptoneurosomes (SYNs) were prepared from whole hippocampal tissue as previously described (Sosanya, Huang et al. 2013, Ewin, Morgan et al. 2019). Briefly, whole hippocampal tissue was homogenized in buffer (50 mM Tris, pH 7.35; protease and phosphatase inhibitors (Halt, Thermofisher). Homogenates were sequentially filtered through 100 μm and 5 μm filters to produce SYNs (Quinlan, Philpot et al. 1999, Niere, Namjoshi et al. 2016). SYNs were centrifuged (14,000g; 20 min; 4°C) to obtain a pellet that was solubilized in RIPA buffer (150 mM NaCl; 10 mM Tris, pH 7.4; 0.1% SDS; 1% Triton X-100; 1% deoxycholate 5 mM EDTA; Halt). Samples were then centrifuged (14,000g; 20 min; 4°C) and the soluble fraction was removed and used for Western blot analysis as described above.

### Glucose tolerance test

After 9 weeks of ethanol exposure, a glucose tolerance test was performed as previously described (Day, Yang et al. 2019). Briefly, mice were fasted for 4 h and 2.0 g/kg glucose was administered via i.p injection. Blood samples were taken from tail veins and blood glucose was measured at baseline, 15-, 30-, 45-, 60-, 90-, and 120 minutes from glucose injection using a glucometer (Bound Tree Medical Precision XTRA Glucometer; Fisher). Glucose tolerance tests were performed on non-drinking days.

### Plasma glucose and lactate measurements

Plasma was collected during euthanasia, as described above. Glucose and lactate concentrations were measured using the YSI 2900 analyzer (YSI incorporated) per the manufacturer’s instructions as previously described (Macauley, Stanley et al. 2015). Data is represented by means ± SEM.

### Insulin ELISA

Plasma was collected during euthanasia, as described above, and insulin was measured by ELISA (Alpco; 80-INSMSU-E10) according to manufacturer’s instructions (Stanley, Macauley et al. 2016). Homeostasis model assessment of insulin resistance scores were then calculated (HOMA-IR = plasma glucose [mmol/L] x plasma insulin [U/mL]/22.5).

### Open Field Assay (OFA)

The open field assay was performed as described previously (Ewin, Morgan et al. 2019). Briefly, mice were placed in the center of a plexiglass chamber (40cm x 40cm x 30cm) equipped with Omnitech Superflex Sensors (Omnitech Electronics, Inc). This box uses arrays of infrared photodectectors located at regular intervals along each wall of the chamber. The chamber walls were solid and were contained within sound-attenuating boxes with a 15-watt light bulb to illuminate the arena. Exploratory activity was measured for 15 minutes and quantified as locomotor activity and % time spent in the central zone. OFA activity was assessed at baseline when mice were ∼3 months-old, and again after 3 weeks ethanol exposure when mice were ∼6 months-old.

### Light/Dark Assay (LD)

The light/dark box test was conducted as previously described (Miller, Piasecki et al. 2011). Control and APP/PS1 mice were placed into a polycarbonate box (40 cm x 40 cm) with two equally sized regions. One region was dark and concealed, while the other was open and light. A 10 cm opening allowed free movement between both regions. Mice were monitored for five minutes. Latency to enter the light side, number of light-side entries, and total time spent in the light-side of the box were recorded with EthoVision XT tracking software. Increased reluctance to venture into the light, uncovered, side was interpreted as anxiety-related behavior. LD activity was assessed at baseline when mice were ∼3 months-old, and again after 3 weeks of ethanol exposure when mice were ∼6 months-old.

### Marble Burying

The marble burying test was performed as previously described (Amodeo, Jones et al. 2012). Control and APP/PS1 mice were brought into a novel environment and habituated for one hour before behavioral testing. Mice were placed in a cage (19.56 cm x 30.91 cm x 13.34 cm) containing 12 marbles (13 mm diameter) on corncob bedding (5 cm depth). Mice were allowed to freely move within the cage for 30 minutes. Following the 30-minute period, mice were removed from the cage and returned to their respective home cages. Images of each cage were recorded, and the number of marbles were counted. A marble was considered buried when >75% of the object was covered by bedding.

### Object location memory task (OLM)

Object location memory task was conducted as previously described (Day, Yang et al. 2019). Mice were habituated to an opaque plastic chamber (40cm x 40cm) with visible spatial cues for 10 min. After 24 h, mice were returned to the chamber with two identical objects and were allowed to freely explore and interact with the objects for 10 min. Twenty-four hours later, mice were returned to the chamber again, where one of the two objects had been relocated to an adjacent position. Changes in objects and locations were randomized and counterbalanced. Time spent with each object was measured and calculated as a percentage of the total object interaction time. Relocated object preference of ∼50% indicates memory impairments. Time with objects was measured both manually and with EthoVision XT tracking software. Mice with a total object interaction time of <5 s were excluded from analysis. Data collection and analysis were performed blinded to condition.

### Nest Building

Nest building behavior was assessed as previously described (Deacon 2006). 24 hours following the last day of EtOH treatment during the dark cycle, control and APP/PS1 mice were provided fresh nesting material (a paper Bed-r’Nest (TheAndersons) and cotton nestlet (Ancare) in their home cages. At the beginning of the light cycle, photos of the nests were recorded and rated on a 1-5 scale by two blinded analysts. A score of 1 was considered a completely unconstructed nest, while a 5 was considered a completed nest that integrated all available materials.

### In vivo microdialysis

To determine how ethanol directly affects ISF Aβ, a separate cohort of 3 month-old APP/PS1 mice was exposed to a single intoxicating dose of ethanol (3.0 g/kg, 15% w/v, i.p). Hippocampal ISF was continuously collected before and after ethanol exposure using in vivo microdialysis as previously described (Macauley, Stanley et al. 2015). Five days prior to acute ethanol exposure, guide cannulas (BASi) were stereotaxically implanted into the hippocampus (from bregma, A/P: −3.1 mm; M/L: −2.5 mm; D/V: −1.2 mm; at 12° angle) and secured into place with dental cement. One day prior to ethanol, 3 month-old APP/PS1 mice were transferred to sampling cages (Bioanalytical Systems). Microdialysis probes (2 mm; 38 kDa molecular weight cut off; BR-style; BASi) were inserted into the guide cannula, connected to a syringe pump and infused with 0.15% bovine serum albumin (BSA, Sigma) in artificial cerebrospinal fluid (aCSF; 1.3mM CaCl2, 1.2mM MgSO4, 3mM KCl, 0.4mM KH2PO4, 25mM NaHCO3 and 122mM NaCl; pH=7.35) at a flow rate of 1 μL/min. Hippocampal ISF was collected hourly, beginning in the early afternoon. Approximately 24 hours later, mice were administered 3.0 g/kg ethanol via i.p. injection from a 15% ethanol (w/v; in 0.9% saline) and ISF was collected for another 24 hours.

### ISF glucose and ethanol measurements

ISF glucose and ethanol concentrations were measured in each ISF sample from 3-month-old APP/PS1 mice (n=4) using the YSI 2900 analyzer (YSI incorporated) per the manufacturer’s instructions.

### Aβ40 ELISA

ISF samples from 3-month-old APP/PS1 mice (n=5) collected from in vivo microdialysis experiments were analyzed for Aβ40 using sandwich ELISAs as previously described (Bero, Yan et\ al. 2011, Roh 2012, Macauley, Stanley et al. 2015). Briefly, Aβ40 was quantified using monoclonal capture antibodies (a generous gift from Dr. David Holtzman, Washington University) targeted against amino acids 33-40 (HJ2). For detection, a biotinylated monoclonal antibody against the central domain amino acids 13-28 (HJ5.1B) was used, followed by streptavidin-poly-HRP-40. The assay was developed using Super Slow TMB (Sigma). Plates were read on a Bio-Tek Synergy 2 plate reader at 650 nm.

### Statistical Analysis

Statistical analyses were performed with GraphPad Prism 5.0 (GraphPad Software, Inc., San Diego, CA). Two-way repeated-measures ANOVAs were used to analyze differences between-subject factors (genotype and time or genotype and alcohol exposure) and *post hoc* analyses (Šídák’s multiple comparisons) were performed for assessing specific group comparisons. Two-way ANOVAs were employed for all other statistical analyses using Tukey’s HSD test for all post hoc analyses. The level of statistical significance was set at *p* ≤ 0.05. Data were expressed as means ±SEM.

## Results

### APP/PS1 mice consume more ethanol than wildtype mice

Ethanol-drinking behavior was characterized in APP/PS1 mice since an earlier study reported an initial increase in ethanol consumption in 3xTg-AD mice (Hoffman, Faccidomo et al. 2019). First, cumulative differences in ethanol consumption and preference were assessed in APP/PS1 and wildtype mice. Over the course of the 10-week study, APP/PS1 mice consumed more ethanol than wildtype mice (Figure 1b, p=0.0047). APP/PS1 mice consumed greater amounts of ethanol early in the study, which may contribute to the differences in overall consumption (Figure 1b). Interestingly, APP/PS1 and wildtype mice displayed a similar ethanol preference over the course of the study (Figure 1c). Ethanol-exposed wildtype and APP/PS1 mice consumed similar amounts of water throughout the course of the study (Figure 1d). However, H_2_O-exposed APP/PS1 mice consistently consumed greater amounts of H_2_O each week, relative to H_2_O-exposed wildtype mice (Figure 1e, p < 0.0001). This resulted in an overall increase in water consumption over the entire 10-week experiment. (Figure 1e, p < 0.0001). Together, these data indicate that APP/PS1 mice drink more ethanol and water relative to wildtype mice.

### Ethanol treatment promotes neurodegeneration in APP/PS1 mice

Neurodegeneration is a major component of AD pathology and AUD (Jack, Bennett et al. 2016, Rehm, Hasan et al. 2019). Therefore, brain atrophy was measured in APP/PS1 and wildtype mice after the 10-week ethanol self-administration regimen. APP/PS1 mice had decreased brain mass compared to wildtype mice; an effect that was exacerbated by ethanol consumption (Figure 2a). This demonstrates a genotype x ethanol interaction that was not observed in the wildtype mice. Interestingly, there were no differences in cortical thickness (Figure 2b) or in hippocampal area (Figure 2c), suggesting that other brain regions were the source of the ethanol-induced brain atrophy.

**Figure 2:**
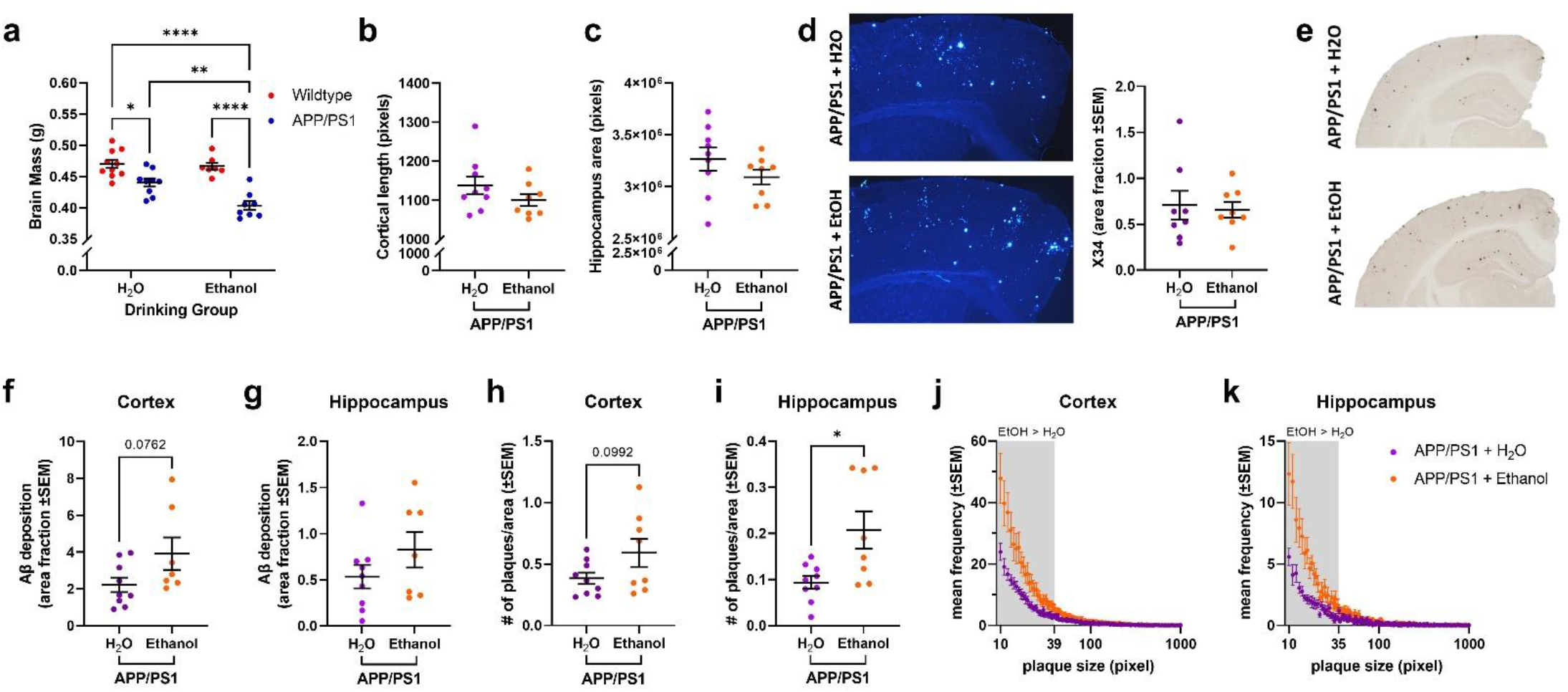
Ethanol exposure increases brain atrophy and amyloid pathology in APP/PS1 mice. a) Brain atrophy was increased in H2O-exposed APP/PS1 mice (p<0.05), an effect that was exacerbated in ethanol-exposed APP/PS1 mice (p<0.01). b) Cortical thickness was comparable between H2O- and ethanol-exposed APP/PS1 mice. c) Hippocampal volume was comparable between H2O- and ethanol-exposed APP/PS1 mice. d) Representative images of X34 staining in cortex of H2O- and ethanol-exposed APP/PS1 mice. There were no differences in X34+ amyloid plaques in the cortex of H2O- and ethanol-exposed APP/PS1 mice. e) Representative images of Aβ deposition in the cortex and hippocampus of H2O- and ethanol-exposed APP/PS1 mice. f) Ethanol-exposed APP/PS1 mice showed a trend towards increased Aβ deposition in the cortex compared to H2O-exposed APP/PS1 mice (p=0.0762). g) No change in Aβ deposition in the hippocampus of H2O- and ethanol-exposed APP/PS1 mice. h) Ethanol-treated APP/PS1 mice had a trend towards increased cortical plaque number compared to H2O-exposed APP/PS1 mice (p=0.0992). i) Ethanol-exposed APP/PS1 mice had increased hippocampal plaque number compared to H2O-exposed APP/PS1 mice (p<0.05). j) Frequency distribution of cortical amyloid plaque size (in pixels). Ethanol-exposed APP/PS1 mice had a greater number of smaller plaques in the cortex compared to H2O-exposed APP/PS1 mice. k) Frequency distribution of hippocampal amyloid plaque size (in pixels). Ethanol-exposed APP/PS1 mice had a greater number of smaller plaques in the hippocampus compared to H2O-exposed APP/PS1 mice. *p<0.05, **p<0.01, ****p<0.0001

### Ethanol treatment increases the frequency of smaller amyloid plaques

Previous studies demonstrate that chronic ethanol consumption at high levels exacerbates AD-like pathology (Huang, Yu et al. 2018, Hoffman, Faccidomo et al. 2019). Therefore, Aβ pathology was quantified following the 10 week moderate drinking paradigm. Quantification of Aβ deposition and amyloid plaques was performed in H_2_O- and ethanol-exposed APP/PS1 mice using HJ3.4B and X34 staining, respectively (Figure 2d-e). While ethanol exposure had no effect on the percent area covered by Aβ deposition or amyloid plaques (Figure 2d), there was a trend towards increased Aβ deposition in the cortex (Figure 2f, p=0.0762). Interestingly, increased plaque number was observed in the hippocampus (Figure 2i, p<0.05) while a trend towards increased plaque number was observed in the cortex (Figure 2h, p = 0.0992). This change in plaque number was associated with a shift in plaque size. Thus, ethanol self-administration was associated with a greater number of smaller plaques in the cortex and hippocampus (Figure 2j-k). This demonstrates that a moderate ethanol-drinking paradigm may promote Aβ pathology by generating smaller plaques. These findings may represent an intermediate stage of plaque proliferation. An ethanol regimen that promotes greater amounts of daily ethanol consumption (Huang, Yu et al. 2018, Hoffman, Faccidomo et al. 2019) or one that runs for a longer duration may induce greater plaque proliferation. Additional studies are needed to assess whether these changes in plaque distribution will lead to increased Aβ pathology over time.

### Ethanol exposure does not alter APP protein levels or metabolism

It is unclear whether the differences in plaque size and number in ethanol-exposed APP/PS1 mice were due to changes in APP expression, APP processing, or Aβ proteostasis. Therefore, APP expression, APP processing, and Aβ degrading enzymes, such as insulin degrading enzyme (IDE), were quantified in this study. APP expression and APP processing (e.g. CTF-β v CTF-α) were comparable between H_2_O- and ethanol-exposed APP/PS1 mice (Figure 3a-c), suggesting moderate ethanol exposure does not affect the amyloidogenic processing of APP. β-secretase (BACE-1), α-secretase (ADAM-10), and insulin degrading enzyme (IDE) expression were similar between H_2_O- and ethanol-exposed APP/PS1 mice (Figures 3d-f). This suggests that enzymes responsible for APP metabolism and Aβ degradation were unaffected by ethanol exposure. Thus, a moderate drinking paradigm does not affect APP levels, amyloidogenic processing of APP, or Aβ degradation.

**Figure 3:**
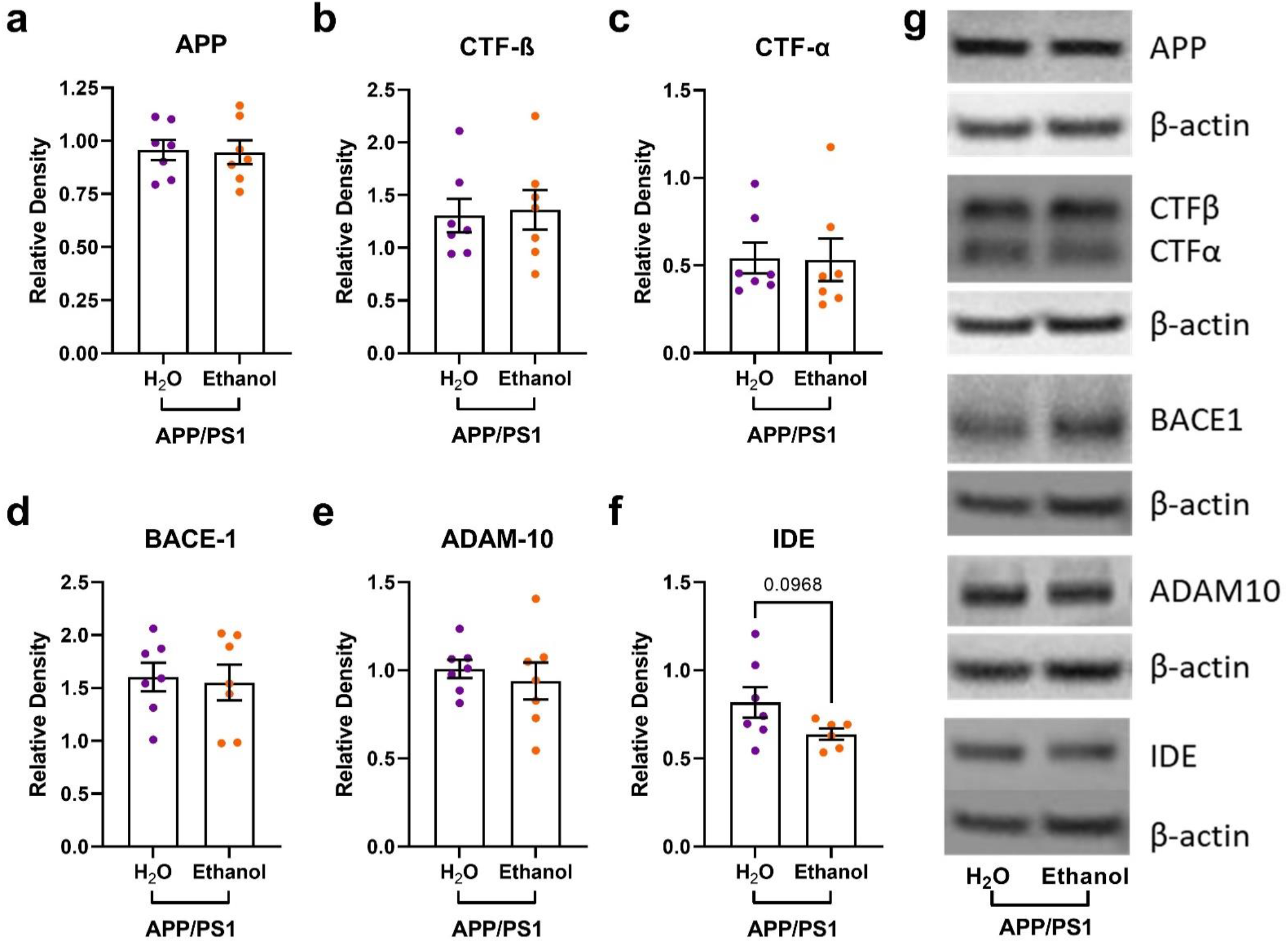
Moderate ethanol drinking does not alter cortical APP metabolism in APP/PS1 mice. a) There were no differences in cortical APP levels between H2O- and ethanol-treated APP/PS1 mice; b) There were no differences in cortical CTF-β levels between H2O- and ethanol-treated APP/PS1 mice; c) There were no differences in cortical CTF-α levels between H2O- and ethanol-treated APP/PS1 mice; d) There were no differences in cortical BACE-1 expression between H2O- and ethanol-treated APP/PS1 mice; e) There were no differences in cortical ADAM-10 levels between H2O- and ethanol-treated APP/PS1 mice; f) There was a trend towards decreased IDE expression between H2O- and ethanol-treated APP/PS1 mice (unpaired t-test, p = 0.0968); g) Representative gels from Western blot experiments.

### Ethanol exposure alters NMDA and GABA_A_ receptor gene expression

In rodents, ethanol directly modulates neuronal excitability and inhibition during ethanol exposure and withdrawal. N-methyl-D-aspartate and γ-aminobutyric acid A receptors (NMDARs and GABA_A_Rs) play important roles in mediating excitability and inhibition during ethanol exposure and withdrawal. NMDAR and GABA_A_R subunit expression is altered in AUD patients and in rodent models after chronic ethanol exposure (Roberto, Bajo et al. 2006, Farris and Mayfield 2014, Gruol, Huitron-Resendiz et al. 2018). Therefore, expression levels for NMDAR (e.g.*Grin2a*, *Grin2b*) and GABA_A_R (e.g.*Gabra5)* were assessed in H_2_O- and ethanol-exposed APP/PS1 and wildtype mice. While no differences in *Grin2a* levels were observed (Figure 4a), *Grin2b* mRNA levels were elevated in ethanol-exposed APP/PS1 mice (Figure 4b, p = 0.0319), suggesting increased NMDA receptor expression in this group. While *Gabra5* expression was elevated in H_2_O-exposed APP/PS1 mice compared to H_2_O-exposed wildtype (Figure 4c, p = 0.0388), this increase was lost in ethanol-exposed APP/PS1 mice (Figure 4c, p = 0.0687). This suggests ethanol decreased GABA_A_ receptor expression specifically in APP/PS1 mice. Next, synaptoneurosomes were isolated from the hippocampus and analyzed via Western blot to explore changes in NMDAR (e.g. GluN2A, GluN2B) and GABA_A_R (e.g. GABA_A_R α5) subunit levels occurring at the synapse. In synaptoneurosomes, GluN2A and GluN2B levels were similar between groups (Figure 4d-e). However, synaptic GABA_A_R α5 levels increased in ethanol-exposed wildtype mice, but this effect was not present in ethanol-exposed APP/SP1 mice (Figure 4f). Together, these results suggest that ethanol-exposure differentially affects NMDA and GABA_A_ receptor expression, levels, and potentially, trafficking to the synapse in APP/PS1 mice compared to wildtype.

**Figure 4:**
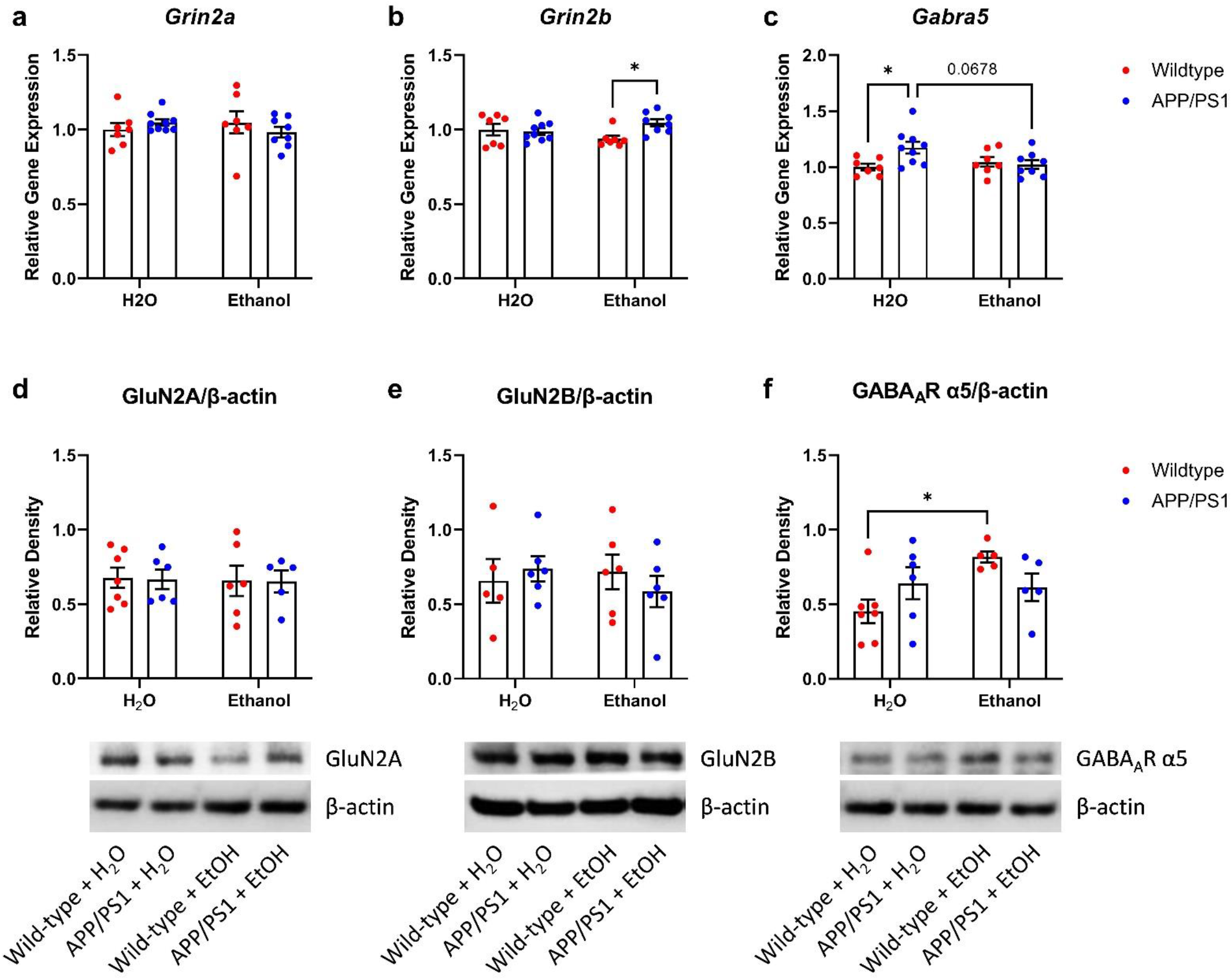
Moderate drinking differentially alters NMDA and GABAA receptors in the cortex and hippocampus of APP/PS1 mice. a) Ethanol treatment did not alter cortical *Grin2a* expression in wildtype or APP/PS1 mice. b) Ethanol-treated APP/PS1 mice had higher cortical *Grin2b* expression compared to EtOH-treated wildtype mice. 2-way ANOVA revealed a significant treatment x genotype interaction (p = 0.0319). c) H2O-treated APP/PS1 mice showed increased cortical *Gabra5* expression compared to H2O-exposed wildtype mice (p<0.05). This effect was lost in EtOH-exposed APP/PS1 *Gabra5* mRNA levels. 2-way ANOVA revealed a significant treatment x genotype interaction (p = 0.0249) and a trend in genotype effects (p = 0.0723). d) Synaptic GluN2A levels was unaltered in the hippocampus of H2O- or EtOH-treated wildtype or APP/PS1 mice. e) Synaptic GluN2B levels was unaltered in the hippocampus of H2O- or EtOH-treated wildtype or APP/PS1 mice. f) Ethanol-treated wildtype mice showed increased synaptic GABAAR α5 subunit levels compared to H2O-treated wildtype mice. Ethanol treatment had no effect on GABAAR α5 subunit levels in APP/PS1 mice. 2-way ANOVA revealed a significant treatment x genotype effect (p = 0.0347) and a trend in treatment effects (p = 0.0644). *p<0.05

### Ethanol exposure dysregulates diurnal food consumption in APP/PS1 mice

AD and AUD are both characterized by disruptions in circadian rhythms (Carroll and Macauley 2019, Koob and Colrain 2020). Thus, food consumption was measured every 12 hours across the diurnal cycle on drinking days for four weeks, starting at 7 months of age (Figure 5a-c). Since mice are nocturnal, they consume the majority of their food during their dark cycle. While food consumption was highest at night for all groups, ethanol-drinking APP/PS1 mice consumed less food during this period than all other groups (Figure 5b). Additionally, ethanol-drinking APP/PS1 mice consumed more food during their light cycle, when mice normally spend more time sleeping (Figure 5c). Interestingly, this effect was not observed in ethanol-exposed wildtype mice (Figure 5b-c). Total food consumption across the 24-hour day was comparable between H_2_O-exposed APP/PS1 and wildtype mice (Figure 5b-c). This diurnal misalignment with food intake suggests that chronic ethanol consumption may disrupt sleep, specifically in mice with Aβ overexpression.

**Figure 5:**
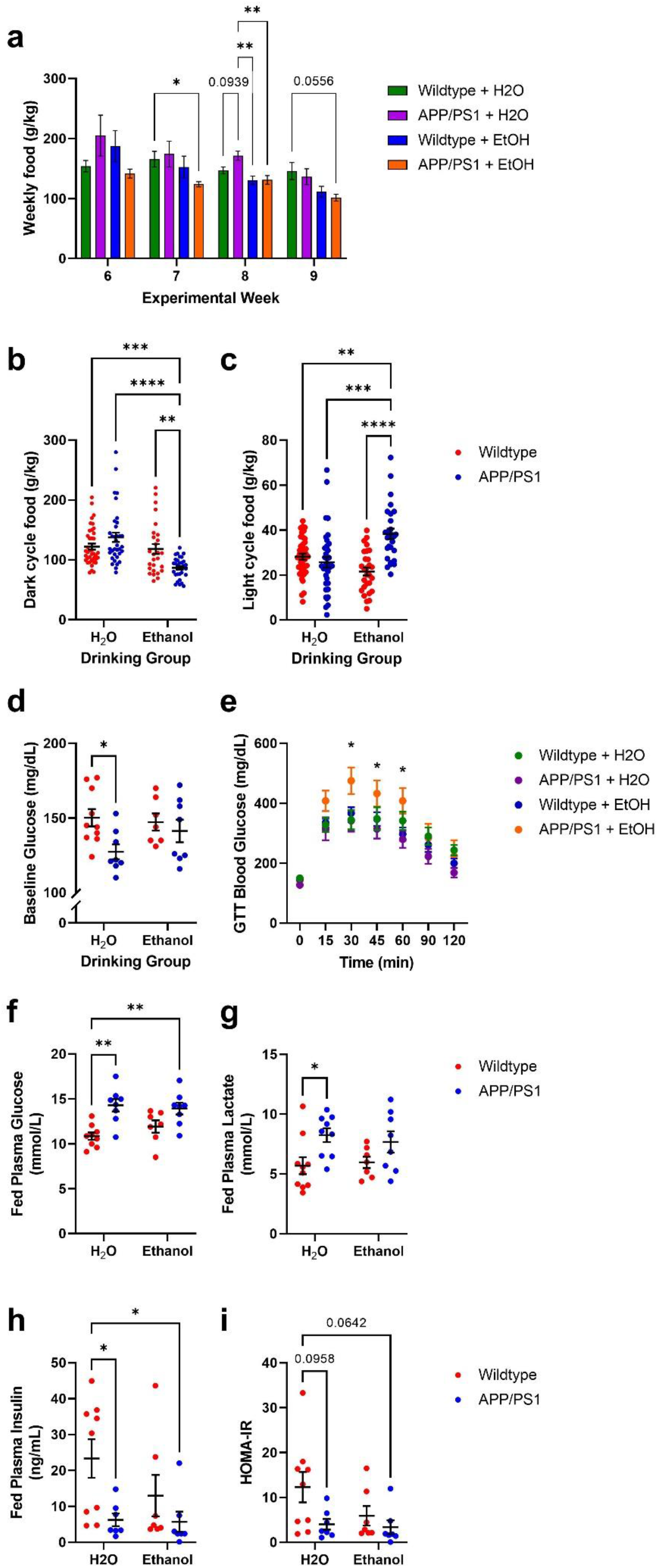
Ethanol exposure dysregulates diurnal feeding behavior and peripheral metabolism in APP/PS1 mice. a) Decreased weekly food consumption in APP/PS1 with or without ethanol treatment during experimental weeks 6-9. b) Ethanol-treated APP/PS1 mice showed decreased food consumption during the dark period. Two-way ANOVA revealed significant differences between drinking group (p<0.0001) and genotype x drinking group (p=0.0003). c) Ethanol-treated APP/PS1 mice showed increased food consumption during light cycle. 2-way ANOVA revealed significant interactions of genotype (p=0.0005) and genotype x drinking group (p<0.0001). d) H2O-treated APP/PS1 mice showed lower fasted blood glucose concentrations prior to glucose tolerance test. 2-way ANOVA revealed significant genotype effect (p = 0.0273). e) Ethanol-treated APP/PS1 mice displayed glucose intolerance during glucose tolerance test. Two-way ANOVA revealed significance over time (p<0.0109), and time x group (p<0.0001). Tukey’s multiple comparisons revealed that EtOH-treated APP/PS1 mice had significantly higher blood glucose concentrations at 30-, 45-, and 60-minutes post-glucose injection. f) H2O- and EtOH-treated APP/PS1 mice had higher plasma glucose at the terminal timepoint of death. 2-way ANOVA revealed genotype effect (p = 0.0001). g) H2O-treated APP/PS1 mice had higher plasma lactate levels at the terminal time point compared to H2O-treated wildtype mice. 2-way ANOVA revealed a genotype effect (p = 0.0049). h) H2O- and EtOH-exposed APP/PS1 mice had decreased fed insulin levels at the terminal timepoint, compared to H2O-treated wildtype mice. 2-way ANOVA revealed a genotype effect (p = 0.0123). i) Calculated HOMA-IR values showed a trend trended towards decreased in H2O- and ethanol-exposed APP/PS1 mice. 2-way ANOVA revealed a genotype effect (p=0.0381). *p<0.05, **p<0.01, ****p<0.0001

### Chronic ethanol exposure alters glucose homeostasis in APP/PS1 mice

Both AUD and AD are associated with metabolic impairment and impaired glucose homeostasis (Macauley, Stanley et al. 2015). Therefore, alterations in glucose metabolism were assessed in 7.5 month-old mice after 9 weeks of ethanol exposure. Fasted blood glucose levels were lower in H_2_O-exposed APP/PS1 mice compared to wildtype mice (Figure 5d); however, this trend was reversed with ethanol exposure. In fact, fed plasma glucose levels were elevated in H_2_O- and ethanol exposed-APP/PS1 mice compared to wildtype mice, suggesting Aβ pathology differentially affects peripheral metabolism (Figure 5f). Interestingly glucose intolerance was only observed in ethanol-exposed APP/PS1 mice during a glucose tolerance test, suggesting ethanol exposure exacerbates metabolic dysfunction and insulin resistance in APP/PS1 mice (Figure 5e). In addition to changes in fed glucose levels, H_2_O-exposed APP/PS1 mice had elevated fed lactate (Figure 5g). Both H_2_O-and ethanol-exposed APP/PS1 mice had decreased fed insulin levels at a terminal timepoint (Figure 5h). They also showed a trend towards increased insulin resistance as measured by HOMA-IR (Figure 5i). Given these differences in peripheral metabolism, body weights were measured for all groups at the beginning and end of the study. At the terminal timepoint, no differences in body weights were observed in this study (data not shown). Taken together these data indicate that moderate levels of chronic ethanol drinking induces metabolic dysfunction specifically in APP/PS1 mice.

### Chronic ethanol consumption alters anxiety- and dementia-related behaviors in APP/PS1 mice

Lower CSF Aβ42, which corresponds to increased amyloid plaques in the brain, was associated with increased anxiety in individuals over the age of 50 (Krell-Roesch, Rakusa et al. 2022). Chronic ethanol exposure can also lead to increased anxiety-related behaviors and deficits in cognition. Here, anxiety-related behaviors were measured using OFA and LD assays at baseline and after 3 weeks of ethanol exposure. Marble-burying, OLM, and nest building tasks were only performed after ethanol exposure. At baseline, APP/PS1 mice showed a trend towards increased locomotor activity during the OFA (Figure 6a, p = 0.0668), but not in the percent time spent in the center zone (Figure 6b). After 3 weeks of ethanol exposure, there was increased locomotor activity in ethanol-exposed APP/PS1 mice (Figure 6a). Post-hoc tests showed that ethanol-drinking APP/PS1 mice spent more time in the central zone than wildtype mice (Figure 6b). No differences in behavior were observed in LD at baseline, or after 4 weeks of ethanol self-administration (Figure 6c). After 5 weeks of ethanol drinking, mice were tested using the marble burying test, where increased marble burying is used as a measure of anxiety-like behavior. Control APP/PS1 mice buried fewer marbles than wildtype mice, while ethanol-drinking APP/PS1 mice did not (Figure 6d). Ethanol-naïve APP/PS1 mice bury fewer marbles, possibly indicating decreased anxiety-like behaviors or disengagement in the task. Because this phenotype is not seen in ethanol-drinking APP/PS1 mice, this may reflect an increase in anxiety-like behavior during withdrawal. After 7 weeks of ethanol treatment, the OLM task evaluated the effects of ethanol treatment on memory. As expected, H_2_O-exposed wildtype mice spent more time interacting with the relocated object than with the object in the familiar location (Figure 6e, p = 0.0078). Conversely, APP/PS1 mice and ethanol-exposed wildtype mice spent similar amounts of time investigating both objects (Figure 6e). This indicates that impaired memory due to Aβ pathology was not exacerbated by ethanol exposure. However, this may be due to a ceiling effect as APP/PS1 control mice exhibit maximal memory impairment on this assay. Lastly, APP/PS1 mice showed deficits in nest building, which was exacerbated by ethanol exposure (Figure 6f). Conversely, ethanol treatment had no effect on nest building behavior in wildtype mice. This further demonstrates that ethanol exposure exacerbates anxiety and AD-related behavioral deficits.

**Figure 6:**
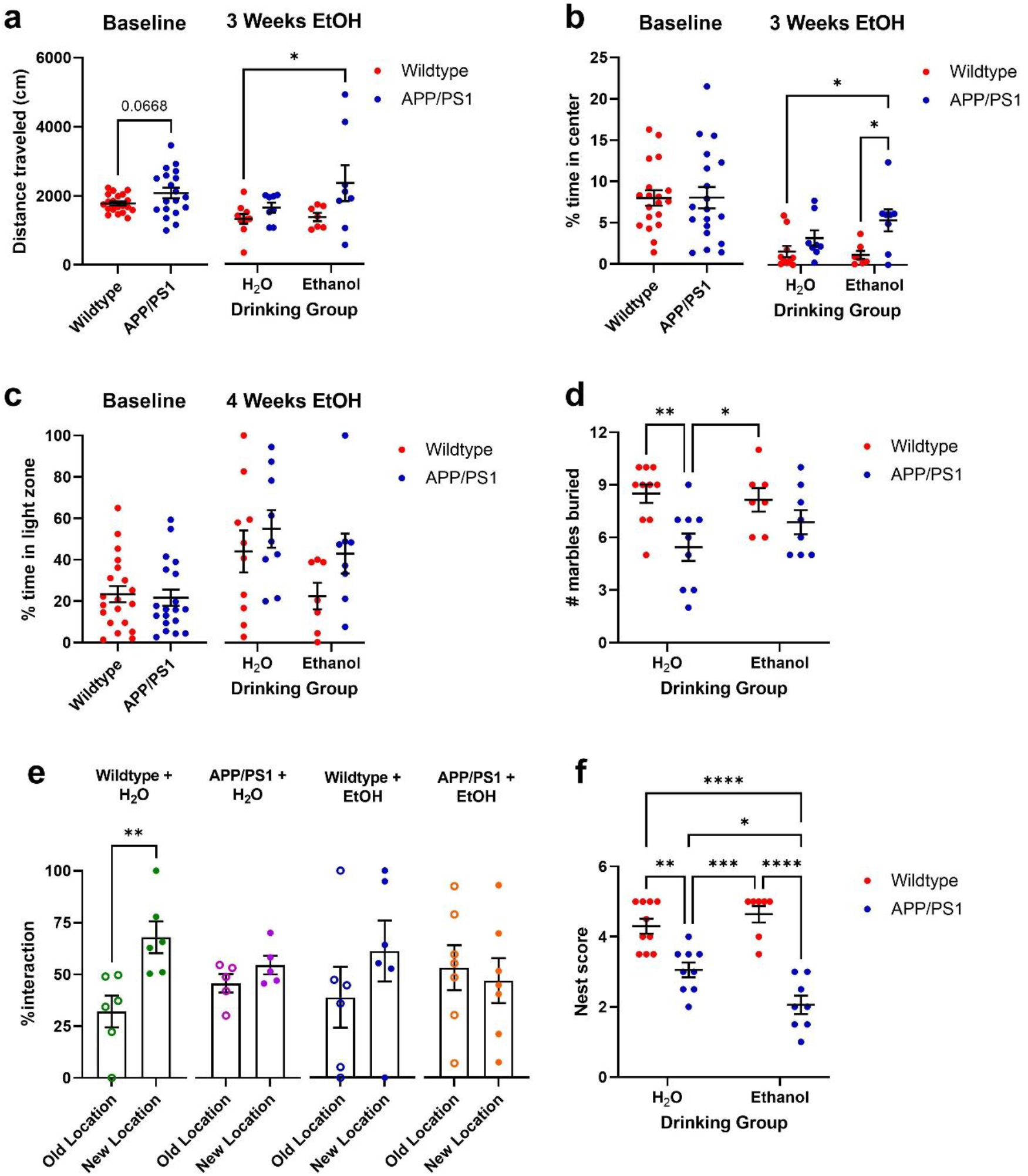
Chronic ethanol consumption alters anxiety-related and dementia-related behaviors in APP/PS1 mice. a) At baseline, APP/PS1 mice showed a trend towards increased locomotor activity during the OFA (unpaired t-test, p = 0.0668). After 3 weeks of ethanol exposure APP/PS1 mice showed more locomotor activity than other groups. b) There were no differences in the % time spent in the center zone at baseline. After 3 weeks of ethanol treatment, APP/PS1 mice spent more time in central zone than wildtype controls. c) Mice exhibited no differences spent in the light zone in the LD box at baseline or following treatment. d) H_2_O-treated APP/PS1 mice buried more marbles than wildtype controls. e) H_2_O-treated wildtype spent significantly more time interacting with the relocated object than with the object in the familiar location (unpaired t-test, p = 0.0078), while other groups spent similar amounts of time interacting with both objects. f) APP/PS1 mice made poorer nests compared to wildtype mice. 2-way ANOVA revealed differences in nest building scores between groups after 9 weeks of EtOH treatment. APP/PS1 mice made poorer nests compared to wildtype mice. This effect was exacerbated by ethanol exposure (2-way ANOVA: genotype: p<0.0001; genotype x drinking group: p=0.0071). *p<0.05, **p<0.01, ***p<0.001, ****p<0.0001

### Acute ethanol exposure directly affects Alzheimer’s-related pathology and brain metabolism

Chronic ethanol exposure altered AD-like pathology and metabolism. Therefore in vivo microdialysis was used to determine whether ethanol directly modulates hippocampal ISF Aβ and glucose levels in unrestrained, unanesthetized APP/PS1 mice (Figure 7a). An acute dose of ethanol (3.0 g/kg, i.p.) led to a rapid increase in ISF ethanol levels (14.79±1.73 mmol/L), which declined over the next 6 hours (Figure 7b-c, Supplementary Table 1), demonstrating ethanol freely crosses the blood brain barrier into hippocampal ISF. Interestingly, ethanol exposure caused bidirectional changes in ISF Aβ levels (Figure 7b). Following acute ethanol exposure, ISF Aβ levels decreased (- 15.82% ± 1.58% below baseline) then rose rapidly and peaked at 6 hours post-ethanol injection (20.07% ± 6.83% above baseline; Figure 7b, Supplementary Table 1). Conversely, an i.p. ethanol injection increased ISF glucose levels in the first hour which returned to baseline over the next 6 hours (Figure 7d, Supplementary Table 1). ISF ethanol and ISF glucose levels were positively correlated during the 6-hour time period that ethanol was present in the brain (Figure 7e, r = 0.6921, p = 0.0068). These data have two important implications. First, they demonstrate that when ISF ethanol is elevated, ISF glucose is elevated as well. This could explain, in part, the glucose intolerance phenotype seen in the chronic study where ethanol led to increased peripheral glucose levels (Figure 5d, f). Second, these data also show that when the hippocampus is exposed to ethanol, less ISF Aβ is released initially. However, when ethanol is cleared from the brain, the brain’s response during withdrawal drives ISF Aβ release and increases the extracellular pool of Aβ. Thus, withdrawal from a single ethanol exposure is sufficient to raise ISF Aβ levels and concurrently dysregulate brain metabolism, two known mediators of amyloid plaque formation.

**Figure 7:**
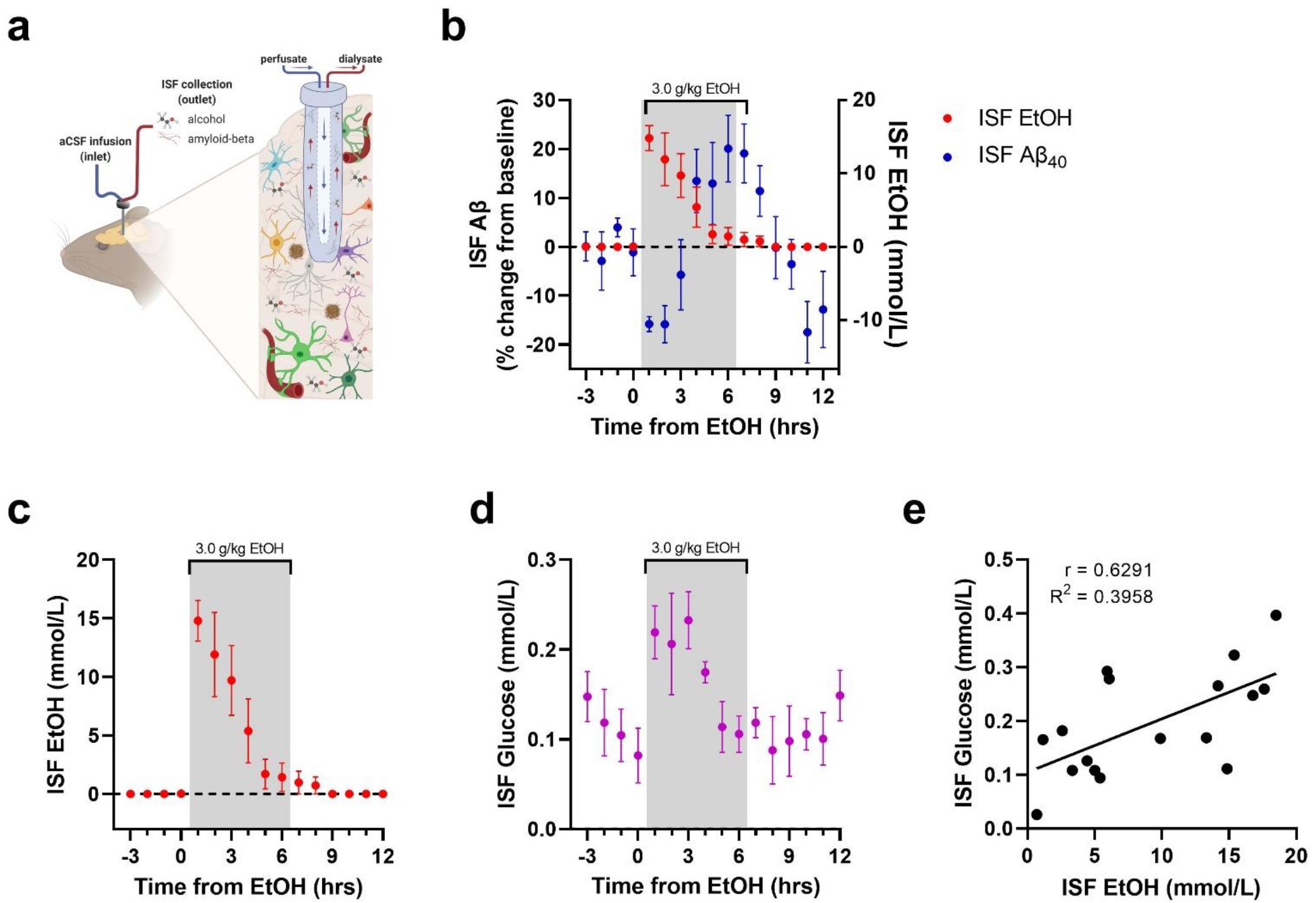
Ethanol acutely modulates ISF Aβ and ISF glucose in APP/PS1 mice. a) Schematic of 38 kDa in vivo microdialysis to sample brain hippocampal interstitial fluid (ISF). b) Ethanol exposure changes ISF Aβ levels (left y-axis; blue) and ISF EtOH (right y-axis; red) in 3 month-old APP/PS1 mice. c) Changes in ISF ethanol in 3 month-old APP/PS1 mice. c) Changes in ISF glucose in 3 month-old APP/PS1 mice. d) ISF ethanol and ISF glucose are significantly correlated while ethanol is present in the brain.

## Discussion

This study found that long-term, moderate ethanol self-administration increases AD-related pathology, alters peripheral metabolism, and exacerbates behavioral deficits in APP/PS1 mice. Ethanol exposure consistently exacerbated phenotypes related to Aβ pathology, neuronal excitatory-inhibitory balance, metabolism, and behavior. This suggests that early changes in Aβ pathology synergize with ethanol exposure to potentiate damage to the brain. We also found that acute ethanol exposure and withdrawal bidirectionally modulated Aβ levels within the hippocampal ISF. To our knowledge, this is the first study to provide evidence that a single exposure of ethanol directly modulates ISF Aβ levels. These findings also build upon existing studies to demonstrate that even at moderate levels of intake, ethanol exposure can worsen AD-related pathology and behavioral impairment.

In humans, amyloid pathology begins to accumulate ∼10-20 years before the onset of clinical symptoms (Jack, Knopman et al. 2010). In APP/PS1 mice, Aβ begins to aggregate into amyloid plaques by 6-9 months of age (Jankowsky, Fadale et al. 2004). Previous studies consistently show that chronic ethanol exposure is sufficient to increase amyloid burden in mouse models of cerebral amyloidosis (Huang, Yu et al. 2018, Hoffman, Faccidomo et al. 2019). Here, APP/PS1 mice showed signs of brain atrophy, as measured by decreased brain mass. Chronic ethanol consumption exacerbated this phenotype in APP/PS1 mice while brain mass in wildtype mice was unaffected by ethanol consumption (Figure 2a). This could be due to an interaction with the formation of amyloid plaques or through a mechanism entirely independent from AD-like pathology. Interestingly, AUD can lead to different forms of dementia, including vascular dementia, Parkinson’s disease, Korsakoff syndrome, or alcohol-related dementia – all of which display alcohol-related brain damage, neurodegeneration, and brain atrophy (Hakon, Quattromani et al. 2018). Future studies are needed to tease apart whether the brain atrophy observed in this study was dependent on changes in the Alzheimer’s cascade.

While ethanol had modest effects on the amount of Aβ deposition, there was an increase in the number of plaques in the cortex and hippocampus (Figure 2h-i). Further analysis showed that ethanol exposure increased the number of smaller plaques in both the cortex and hippocampus (Figure 2j-k). This could be due to ethanol increasing the formation of new smaller plaques, or conversely, restricting plaque growth. First, chronic ethanol may exacerbate amyloid pathology by increasing the number of smaller plaques at an age that corresponds to presymptomatic AD. A greater number of smaller plaques could create multiple pro-aggregation sites or plaque seeds, ultimately leading to increased plaque proliferation later in life. Alternatively, the emergence of smaller amyloid plaques could suggest that low-to-moderate ethanol consumption somehow restricts plaque growth, leading to smaller plaques. Interestingly, these changes in plaque number and size distribution did not correspond with changes in APP levels, APP metabolites, or APP/Aβ degrading enzymes (Figure 3). Future studies should explore whether these changes in plaque size and plaque number are a harmful or protective response to moderate alcohol consumption. Nevertheless, the other findings from this study clearly demonstrate an ethanol-associated reduction in brain mass, specifically in APP/PS1 mice. Thus, it is more likely the increased plaque number and reduced plaque size were early aggregatory events that would be potentiated if dose and duration of ethanol exposure was increased.

In vivo microdialysis explored how ethanol modulates ISF Aβ levels in unrestrained and unanesthetized APP/PS1 mice. In this study, ethanol bidirectionally altered ISF Aβ levels. ISF Aβ levels dramatically decreased in response to an acute ethanol exposure then increased during withdrawal (Figure 7b). Ethanol is known to directly modulate neuronal activity by increasing GABA inhibition during exposure and increasing NMDA hyperexcitability during withdrawal (Ariwodola and Weiner 2004, Slawecki, Roth et al. 2006, Weiner and Valenzuela 2006, Cheaha, Sawangjaroen et al. 2014, Ramachandran, Ahmed et al. 2015, Wang, Zhao et al. 2016, Roberto and Varodayan 2017). Because Aβ is released from neurons in an activity-dependent manner (Cirrito, Yamada et al. 2005, Bero, Yan et al. 2011, Verges, Restivo et al. 2011), the changes in ISF Aβ may be due to ethanol-induced changes in excitation during intoxication and withdrawal. In fact, preclinical and postmortem studies demonstrate that the NMDAR subunits, GluN2A and GluN2B, are upregulated in rodents after a chronic ethanol exposure as well as in humans with AUD (Roberto, Schweitzer et al. 2004, Farris and Mayfield 2014). The effects of chronic ethanol on GABA_A_Rs are well-documented (Roberto and Varodayan 2017), and the GABA_A_R α5 subunit is modulated by chronic ethanol in preclinical studies at the gene and protein level (Centanni, Teppen et al. 2014, Gruol, Huitron-Resendiz et al. 2018, Zeng, Xie et al. 2019). Therefore, ethanol-induced inhibition and excitation may drive the activity-dependent production of Aβ, representing an early pathological mechanism by which chronic ethanol exposure drives plaque deposition. In the chronic studies, there were differences found in GluN2B and GABA_A_R α5 mRNA and protein levels in response to moderate ethanol exposure (Figure 4). Interestingly, increases in cortical *Grin2b* mRNA levels between wildtype and APP/PS1 mice were only induced by chronic ethanol exposure. Ethanol drinking had the opposite effect on *Gabra5* expression, decreasing *Gabra5* to wildtype levels (Figure 4b-c). While the effect size was small, these changes corresponded with a trend towards increased Aβ deposition and plaque number in the cortex of ethanol-exposed APP/PS1 mice (Figures 2g, 2i). Furthermore, these modest effects may also be due to the moderate ethanol exposure paradigm. Future studies will explore how higher doses of ethanol alters neuronal excitability and inhibition and potentially drive plaque deposition in APP/PS1 mice.

Chronic ethanol consumption can lead to increased anxiety- and depression-related behaviors, which can be further exacerbated by changes in glucose metabolism (Bouwman, Adriaanse et al. 2010). During weekly withdrawal periods, ethanol drinking APP/PS1 mice exhibited changes in locomotor activity and central zone exploration in the OFA (Figure 6a-b) but not in LD exploration (Figure 6c). Ethanol did not induce these behavioral changes in wildtype mice, which may indicate that APP/PS1 mice are especially sensitive to anxiety-related behaviors. Alternatively, ethanol drinking may exacerbate the hyperactive, impulsive, or compulsive phenotype observed in APP/PS1 mice (Shepherd, May et al. 2021). Conversely, ethanol treatment did not aggravate the deficits in the marble-burying task observed in APP/PS1 controls (Figure 6d). Thus, further tests should be conducted to better understand the behavioral phenotypes reported here.

Ethanol’s effects on cognition are well-documented (Sabia, Elbaz et al. 2014) and the present study indicates that chronic ethanol drinking may be detrimental to long-term memory in wildtype mice. In the object location memory task (OLM), ethanol exposure disrupted memory in wildtype mice but had no effect on APP/PS1 mice (Figure 6e). Future studies should employ more sensitive cognitive tasks to identify the degree to which ethanol affects cognition in APP/PS1 mice. One major deficit commonly observed in patients with mild cognitive impairment is a disruption in self-care behaviors (i.e. cleaning one’s room, showering, etc.) (Hirschfeld, Montgomery et al. 2000). In this study, a major impact of ethanol exposure on self-care-related behaviors was observed, as measured by nest building (Jirkof 2014). Control (H_2_O-drinking) APP/PS1 mice displayed reduced nest-building behaviors, and this deficit was exacerbated in ethanol-exposed APP/PS1 mice (Figure 6f). This outcome further suggests that ethanol-drinking APP/PS1 mice are experiencing earlier onset of negative affective behaviors associated with AD progression. Interestingly, these differences in nest-building behavior were similar to changes in brain mass at the end of the study. While ethanol exposure had no effect on brain mass in wildtype mice, this measure was reduced in control APP/PS1 mice and was further reduced in ethanol-drinking APP/PS1 mice (Figure 2a).

Ethanol-exposed APP/PS1 mice developed an interesting metabolic phenotype in both the acute and chronic studies. In the acute studies, in vivo microdialysis measured how ethanol modulates cerebral glucose metabolism in APP/PS1 mice during ethanol exposure. An i.p. ethanol injection rapidly increased ISF glucose levels (Figure 7d), which correlated with ISF ethanol concentrations in the hippocampus. This change in ISF glucose is sufficient to drive changes in ISF Aβ levels based on previous research that demonstrated hyperglycemia increases ISF Aβ release (Macauley, Stanley et al. 2015). This offers a metabolic explanation for the link between AUD and AD. The chronic studies also reinforced the idea that moderate ethanol consumption alters food intake and peripheral metabolism. First, APP/PS1 mice demonstrated alterations in fed glucose, lactate, and insulin levels as well as insulin resistance when compared to wildtype mice (Figure 5f-i). This suggests Aβ pathology disrupts glucose homeostasis independent of ethanol. However, ethanol exposure caused glucose intolerance, most likely due to hypoinsulinemia, but only in APP/PS1 mice. These changes were not observed in ethanol-exposed wildtype mice, suggesting that Aβ interacts with ethanol to exacerbate changes in metabolism in an AD-specific manner. While metabolic diseases like type-2 diabetes are known to put the brain at risk for AD (Ott, Stolk et al. 1999, Arnold 2018), a growing body of evidence suggests that AD can also exacerbate metabolic dysfunction, glucose intolerance, or insulin resistance. This metabolic dysfunction can be further exacerbated by alcohol. Collectively this supports previous literature showing that alcohol intake leads to dysregulation of peripheral and cerebral metabolism.

Despite glucose intolerance, the APP/PS1 mice did not eat more. However, they did have alterations in diurnal eating patterns with less food consumed during their dark cycle (Figure 5b) and more food during their light cycle (Figure 5c). This diurnal mismatch in feeding behavior suggests that APP/PS1 mice also have altered sleep/wake cycles, which may potentiate their metabolic dysfunction. A bidirectional relationship exists between AD and sleep impairment, where disrupted sleep increases AD risk, and increased Aβ and tau aggregation further disrupts sleep (Carroll and Macauley 2019). AUD is also sufficient to disrupt sleep (Koob and Colrain 2020). Thus, alterations in sleep and diurnal rhythms, like those observed with feeding behavior, may provide one explanation for why AUD increases AD risk. Interestingly, a recent study also demonstrated that nest-building increases during proximity to sleep in mice (Sotelo, Tyan et al. 2022). Although further studies are needed, these findings suggest that chronic ethanol drinking exacerbates disruptions in metabolism and sleep that are frequently observed in AD.

## Conclusions

Contrary to some prior clinical and preclinical findings, these findings demonstrate that chronic intake of moderate amounts of ethanol can exacerbate behavioral and pathological AD-like phenotypes in APP/PS1 mice. Not only does a moderate drinking paradigm lead to a shift in amyloid plaque development, but it may also lead to changes in GABA_A_ receptors that mediate the brain’s E/I balance. Finally, this study demonstrates that a single ethanol exposure bidirectionally alters ISF Aβ levels, possibly reflecting the biphasic effects of acute ethanol on neuronal inhibition and excitability. Independent of changes in the E/I balance, acute and chronic ethanol exposure profoundly impacted peripheral and CNS metabolism, both of which exacerbate AD-related pathology. To conclude, these findings contribute to the growing body of evidence that suggests chronic alcohol consumption may represent an important, modifiable risk factor for AD. Future studies will further characterize the biological mechanisms by which chronic ethanol intake promotes and exacerbates AD-related pathology.

## Author contributions

**SCG, SMD:** Conceptualization, Investigation, Formal Analysis, Writing – Original Draft, Writing – Reviewing and Editing. **CWC, JAS, NIN, HK, WV:** Investigation. **SLM, JLW:** Supervision, Conceptualization, Investigation, Formal Analysis, Writing – Original Draft, Writing – Reviewing and Editing, Funding Acquisition.

## Funding

We would like to acknowledge the following funding sources: T32AA007565 (SCG), T32AA007565 (SMD), K01AG050719 (SLM), R01AG068330 (SLM), BrightFocus Foundation A20201775S (SLM), Charleston Conference on Alzheimer’s disease New Vision Award (SLM), Averill Foundation (SLM), P50AA26117 (JLW)

---

**Supplementary Table 1:** Mean ISF EtOH concentrations and ISF Aβ40 levels in response to acute EtOH injection (3.0 g/kg, Figure 7b-e).

| | ISF EtOH (mmol/L) | | ISF Aβ40 (% BL) | | ISF Glucose (mmol/L) | |
| --- | --- | --- | --- | --- | --- | --- |
| Time (hrs) | Mean | ±SEM | Mean | ±SEM | Mean | ±SEM |
| -3 | 0.00 | 0.00 | 0.10 | 2.99 | 0.15 | 0.03 |
| -2 | 0.00 | 0.00 | -2.90 | 5.94 | 0.12 | 0.04 |
| -1 | 0.00 | 0.00 | 3.92 | 1.92 | 0.10 | 0.03 |
| 0 | 0.03 | 0.03 | -1.12 | 4.79 | 0.08 | 0.03 |
| 1 | 14.79 | 1.73 | -15.82 | 1.58 | 0.22 | 0.03 |
| 2 | 11.91 | 3.60 | -15.83 | 3.79 | 0.21 | 0.06 |
| 3 | 9.71 | 2.99 | -5.74 | 7.20 | 0.23 | 0.03 |
| 4 | 5.38 | 2.73 | 13.43 | 6.50 | 0.17 | 0.01 |
| 5 | 1.71 | 1.28 | 12.92 | 8.41 | 0.11 | 0.03 |
| 6 | 1.43 | 1.21 | 20.07 | 6.83 | 0.11 | 0.02 |
| 7 | 0.98 | 0.98 | 19.09 | 6.01 | 0.12 | 0.02 |
| 8 | 0.72 | 0.72 | 11.41 | 5.19 | 0.09 | 0.04 |
| 9 | 0.00 | 0.00 | -0.19 | 6.34 | 0.10 | 0.04 |
| 10 | 0.00 | 0.00 | -3.55 | 5.10 | 0.11 | 0.02 |
| 11 | 0.00 | 0.00 | -17.49 | 6.28 | 0.10 | 0.03 |
| 12 | 0.00 | 0.00 | -12.82 | 7.78 | 0.15 | 0.03 |

## Notes

### Competing Interest Statement

The authors have declared no competing interest.

